# Exosomes are predominantly loaded with mRNA transcript encoding cytoplasmic proteins and exclude mRNA transcript encoding nuclear proteins

**DOI:** 10.1101/2020.07.29.227223

**Authors:** Shabirul Haque, Sarah R. Vaiselbuh

**Author notes:** Correspondence: Sarah R. Vaiselbuh, MD, MBA, Associate Professor in Pediatrics & Molecular Medicine, Zucker School of Medicine, Assistant Professor, Center for Oncology & Cell Biology, The Feinstein Institute of Medical Research, Director, Pediatric Hematology/Oncology, Children’s Cancer Center, Staten Island University Hospital at Northwell Health. **Communication and portal handling by**: Shabirul Haque, PhD, Feinstein Institute for Medical Research, Northwell Health, Manhasset, New York, and.

## Abstract

Exosomes are nanovesicles (∼30-150 nm diameters) released via an endocytic pathway in almost all mammalian cell types. Exosomes are composed of a lipid bilayer membrane that encloses RNA, miRNA, proteins and DNA. This manuscript unravels how exosome cargo is collected by a highly precise process delineating two separate mRNA transcript entities encoding cytoplasmic and nuclear proteins separately.

Ultracentrifuge isolated exosomes were directly converted into cDNA (Exo-cDNA), by a method developed in our laboratory. Cellular RNA was extracted from each cell line and cDNA was prepared (Cell-cDNA). We amplified mRNA transcripts translating *cytoplasmic* proteins CD10 and CXCR4 and mRNA transcripts translating *nuclear* proteins such as proliferating cell nuclear antigen (PCNA), CREB-BP, activation induced cytidine deaminase (AID), and terminal deoxynucleotidyl transferase (TdT). We amplified all four different mRNA transcripts (PCNA, CREB-BP, AID, and TdT) from *cellular* cDNA but none from *exosomal* cDNA (Exo-cDNA). These findings suggest that exosomes carry mRNA transcripts encoding *cytoplasmic* proteins only but mRNA transcripts encoding *nuclear* proteins could not be detected. This important observation could prove to be crucial for the exosome research community since it sheds light on one of the limitations relating to the use of exosomes as biomarkers in cancer biology and other diseases.

**Graphical Abstract:** 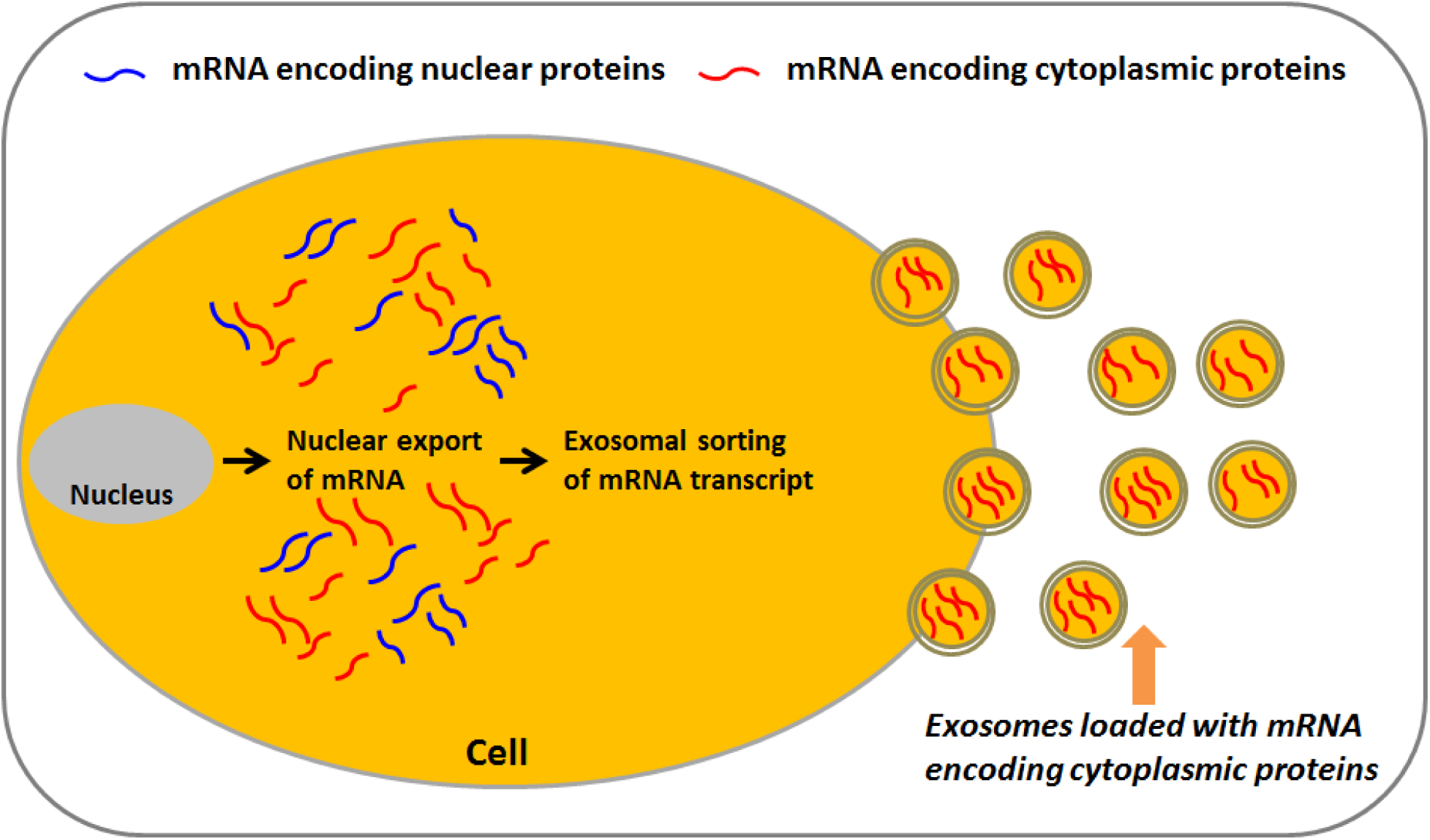

## 1. Introduction

Extracellular vesicles such as exosomes are spherical nanoparticles derived from almost all kind of cells in normal and pathological conditions. The exosome diameter size varies from 30-150 nm, and they originate from the cytoplasm of the cell by inward budding of multivesicular bodies [1]. Exosomal content is rich in proteins, DNA, mRNA and non-coding RNA [2, 3]. Exosomes have been isolated *in vivo* from most bio-fluids such as saliva, serum, urine, breast milk, cerebrospinal fluid, and amniotic fluid [4-7]. In addition, exosomes have been isolated *in vitro* from conditioned medium (CM) of cultured cell lines by ultracentrifugation method [8]. The application of exosomes as biomarkers for disease in cancer research has gained momentum to facilitate early diagnosis, early relapse, risk stratification, and real-time monitoring of the disease in a real time strategy. Exosomes carry the molecular signature of the parent cells. Their major functions are being cell-to-cell communication and exchange of bio-active molecules (such as mRNA and miRNA, DNA, proteins etc.) into target or recipient cells. These exosomal cargo molecules are delivered by a process called ‘exosomal shuttle RNA’ (esRNA) which remains biologically functional in recipient cells [9].

Gene transcription is a multistep and multistage complex process. After mRNA transcription in the nucleus, mRNA transcripts pass through several nuclear modification/process (e.g. 5 prime capping, splicing, and 3 prime ends processing etc.) before being exported to the cytosol. These modifications direct the fate of mRNA transcripts (such as stability, life span, folding etc.) into the cytosol [10]. When processed mRNA transcripts get out of the nucleus they are exposed to the cytosolic milieu in the cytosol. The critical point is, if the mRNA transcripts are poorly processed, they fail to survive and sustain in the cytosolic medium. Several proteins are responsible for nuclear processing of mRNA transcripts, as a result of which many proteins are adhered or anchored onto the mRNA transcripts. These adhered proteins on an mRNA transcript also regulate its export, localization, translation, and stability [10]. It has been established that exosomes collect its entire cargo such as proteins, lipids and coding and non-coding RNAs & DNA directly from the cytoplasmic and/or cytoplasmic membrane derived particles [9]. There are also a few reports which illuminates how different varieties of exosomal RNA species get packaged and loaded in a stoichiometric fashion depending on their source. Typically, serum-derived exosomes contain 30-75% miRNA, 10-20% mRNA, 5-30% rRNA while urine-derived exosomes showed 2-10% miRNA, 5-12% mRNA, 30-60% rRNA [11]. Another study showed that within exosomal RNA, rRNA (28S and 18S fragment) are the most abundant (accounting for ∼97%) and 2% are known RNA (miRNA, mRNA etc.), while 1% are unknown genes [12]. It appears from literature that the knowledge on exosomal RNA content is quite conflicting and limited, and as exosome research is still evolving, investigators are trying to learn about packaging of mRNA transcripts into exosomes.

We hypothesized that selection of mRNA transcripts by the exosomes during exosomal biogenesis is a very precise process. Exosomes selectively entrap mRNA transcripts encoding cytoplasmic proteins, per exclusion of mRNA transcripts that encode nuclear proteins.

## 2. Material and Methods

### 2. 1. Cell lines and reagents

Cell lines of acute lymphatic leukemia (ALL), SUP-B15 (cat# ATCC® CRL-1929 ™), and JM1 cells (cat # ATCC® CRL-10423™) were obtained from ATCC. CL-01 cells obtained (Chiorazzi lab, The Feinstein Institute for Medical research, Northwell Health, Manhasset, NY). Agarose (cat # CA3510-8, Denville Scientific, Inc.). Tris-borate-EDTA buffer (cat # 28355, Thermo Scientific), SYBR Safe (cat # S33102, Invitrogen), 6x loading dye (cat # R0611, Crystalgen). RNase Inhibitor (4 unit/µl) (cat # 1055213, Qiagen), rRNasin Rnase inhibitor (40 U/µl) (Cat # N251B, Promega), RT-Superscript III (200 U/µl) (Cat # 18080-044, Invitrogen), random hexamer primers (cat # SO142, Thermo fisher scientific).

### 2. 2. Cell culture

JM1 cells were expanded and cultured in IMDM supplemented with 0.05 mM β-mercaptoethanol, 10 % FBS, and 1x Penicillin-Streptomycin. SUP-B15 cells were cultured in Iscove’s Modified Dulbecco’s Medium (IMDM), supplemented with 1.5 g/L sodium bicarbonate, 0.05 mM β-mercaptoethanol, 20% fetal bovine serum (not heat inactivated), and 1x Penicillin-Streptomycin. CL-01 cells were expanded and cultured in RPMI-1640 supplemented with 10 % FBS, and 1x Penicillin-Streptomycin. Each cell lines were incubated in a humidified incubator at 37 °C with 5% CO_2_.

### 2. 3. Depletion of exosomes from FBS (Exosome-free FBS)

Fetal bovine serum (FBS) are abundantly loaded with exosomes. FBS was converted into exosome-free FBS by centrifugation and ultracentrifugation method [8]. In short, FBS was subjected to centrifugation for 10 min at 300 xg. Supernatant was collected and centrifuged for another 10 min at 2000 xg. Supernatant was collected again and centrifuged for another 30 min at 10,000 xg. Supernatant was then placed into ultra-centrifugation for overnight at 100,000 xg. FBS supernatant was collected next day and labeled as exo-free FBS. Each centrifugation step was carried out at 4°C [8].

### 2. 4. Exosome isolation from conditioned medium of cell lines

Cell line was cultured in cell culture medium supplemented with Exo-free FBS (FBS depleted with exosomes). Cells (1×10^6^/ml) were plated in 10 ml total volume in a 100 mm tissue culture dish. After 48 hours, CM of each cell line was harvested by centrifugation and filtered through a 0.22 micron filter system to remove cell debris. Filtered CM subjected for exosome isolation by the method of ultracentrifugation.

### 2. 5. Exosome isolation

Exosomes were isolated by the method of centrifugation and followed by ultracentrifugation [8]. Human serum and conditioned medium of cell lines were subjected to centrifugation for 10 min at 300 xg. Collected supernatant was again centrifuged for 10 min at 2000 X g. Supernatant was collected and centrifuged for 30 min at 10,000 xg. Then supernatant was ultra-centrifuged for 120 min at 100,000 xg. Pellets after ultracentrifugation were re-suspended in PBS and ultra-centrifuged for another 120 min at 100,000 xg. Pellets containing exosomes were harvested and reconstituted in around 100-200 μl ice-cold PBS. Each centrifugation and ultra-centrifugation steps were carried out at 4°C. Exosomal protein content was estimated by BCA protein assay kit (Bio-Rad) and stored at -80°C for further usage.

### 2. 6. Direct conversion of exosomes into cDNA (Exo-cDNA)

Exosomes were directly converted into cDNA by following a method published by Haque and Vaiselbuh [13].

### 2. 7. Cellular RNA isolation and cDNA synthesis

Total RNA was extracted from SUP-B15, JM1, and CL-01 cells by the method of Trizol (Invitrogen). RNA concentration and quality was analyzed by NanoDrop ND1000 spectrophotometer. We took cellular RNA (2-5 µg) for cDNA synthesis. We used Oligo-dT primers (Invitrogen), M-MLV reverse transcriptase (Invitrogen) and followed manufacturer’s protocol for cDNA synthesis.

### 2. 8. Gene amplification by two-step PCR

Gene amplification such as β-actin, CD10, CXCR4, PCNA, CREB-BP, AID, and TdT was carried out by two-step PCR [13]. In short, 1^st^ round of PCR was carried out using AccuPrime™ Pfx DNA Polymerase. 2^nd^ round of PCR was carried out using Ampli Taq Gold. Both rounds of PCR are ran under the following conditions: denaturation at 95°C for 30 seconds, annealing at 56°-60°C for 60 seconds, and extension step at 72°C for 60 seconds for 35 cycles. Cell-cDNA or Exo-cDNA (2 µl) were used as template for the first round of PCR. Around 1 – 2 µl of the PCR product of the first round was used as template for the second round of PCR, AmpliTaq Gold was used to a final volume of 30 µl. Brief details of the primers such as gene name, gene bank, sequences, and annealing temperature are mentioned (**Table 1)**.

**Table 1:**
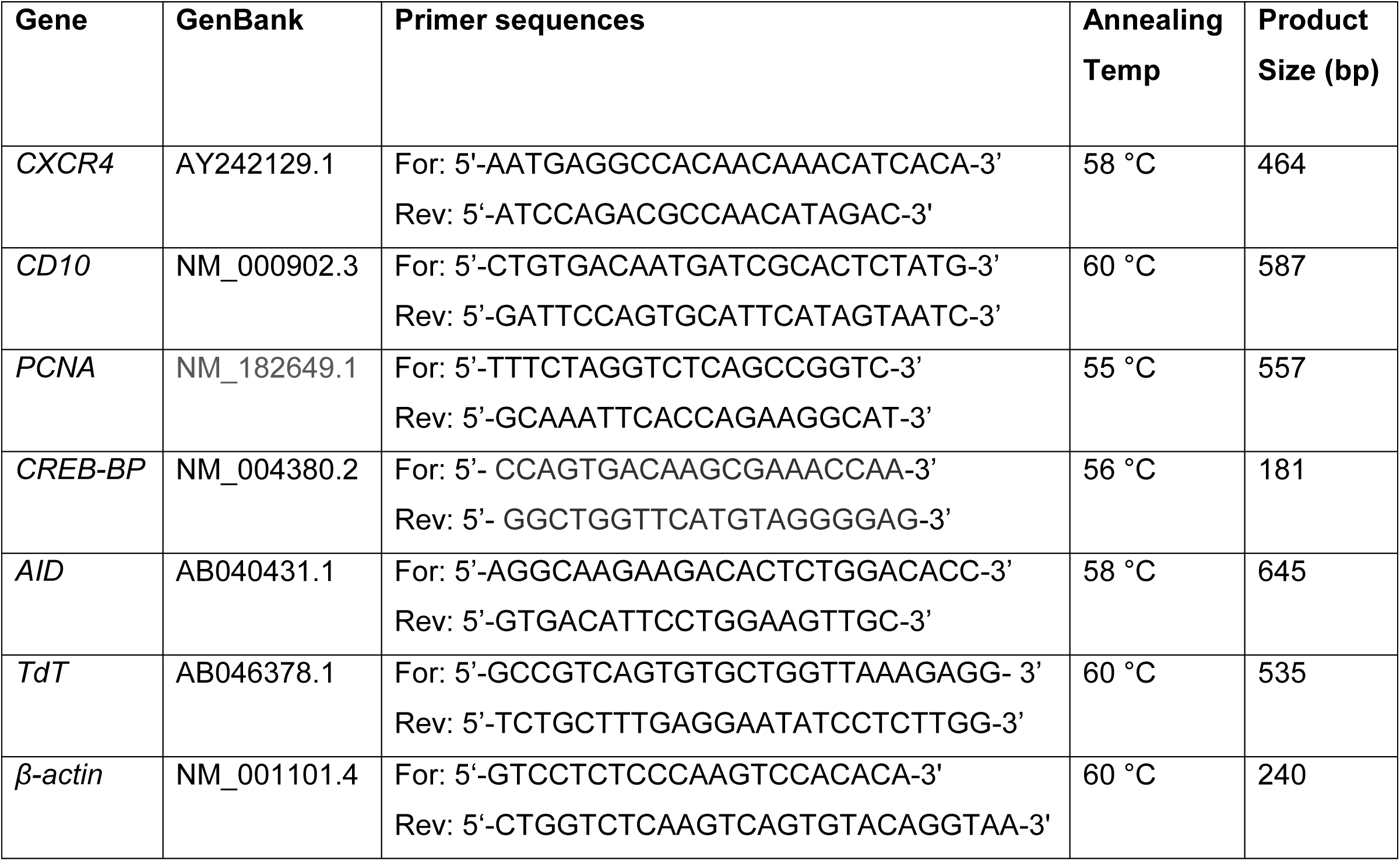
Details of human primers.

### 2. 9. Agarose gel electrophoresis

Briefly, 1.5% agarose was prepared in 1x Tris-borate-EDTA buffer. SYBR Safe was added to the melted agarose gel before polymerization. PCR product (5.0 µl) were mixed with 6x loading dye, mixed product were loaded on to gel and ran for 20-25 minutes at 200 V. The gel was exposed and image were captured by Bio-Rad gel doc system (Bio-Rad, Hercules, California, USA).

## 3. Results

### 3.1. Characterization of exosomes by Nanoparticle Tracking Analysis (NTA)

Exosomes were purified from CM by ultra-centrifugation method. Optimum diameter size of isolated exosomal vesicles was confirmed by Nanoparticle Tracking Analysis (NTA– Malvern) (**Fig. 1 A**). Representative NTA-deduced image of exosomes shown (**Fig. 1 B**). Enrichment of CD63 on exosomes and no expression of calnexin (negative exosomes marker) shown in our published article (13).

**Figure 1.**
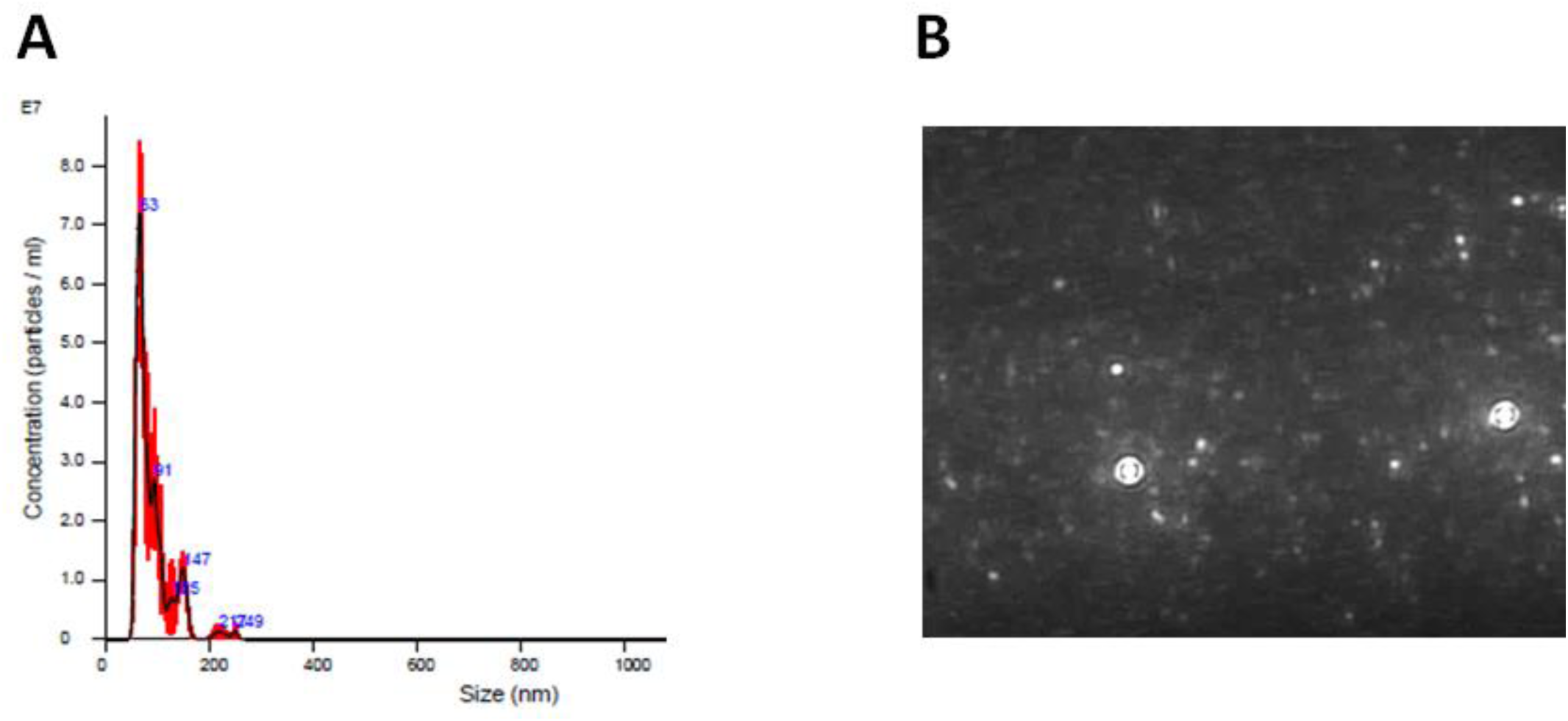
Characterization of exosomes by Nanoparticle Tracking Analysis (NTA). Purified exosomes were passed through NTA system. **A**. Graph shows the heterogeneity of diameter size of the exosomes particle. **B**. Shows representative image of exosomes.

### 3. 2. Amplication of CD10 and CXCR4 from Cell-cDNA/Exo-cDNA

Both the CD10 and CXCR4 proteins are localized within the cytoplasm of the cell. Cell-cDNA and Exo-cDNA from the leukemia cell lines JM1 and SUP-B15 and control normal B-cells CL-01 were used as templates to amplify CD10 and CXCR4 in two-step PCR (**Fig. 2**). We chose CD10 as a marker of the above mentioned leukemia cell lines with CL-01 as negative control. Cellular expression of CD10 was amplified in JM1 and SUP-B15 Cell-cDNA but at exosomal level (Exo-cDNA), CD10 expression could not be detected after a first round of PCR (**Fig. 2 A, upper panel**). After second round of PCR, CD10 expression was detected in both Cell-cDNA and Exo-cDNA (**Fig. 2 A, lower panel**). CXCR4 was amplified by PCR using Cell-cDNA obtained from JM1, SUP-B15, and CL-01 cells but remained undetectable in Exo-cDNA in a first round of PCR (**Fig. 2 B, lower panel**). A second round of PCR showed amplification of CXCR4 expression in both Cell-cDNA and Exo-cDNA (**Fig. 2 B, lower panel**). Endogenous control, β-actin was considered internal control for quality confirmation of Cell-cDNA and Exo-cDNA.

**Figure 2.**
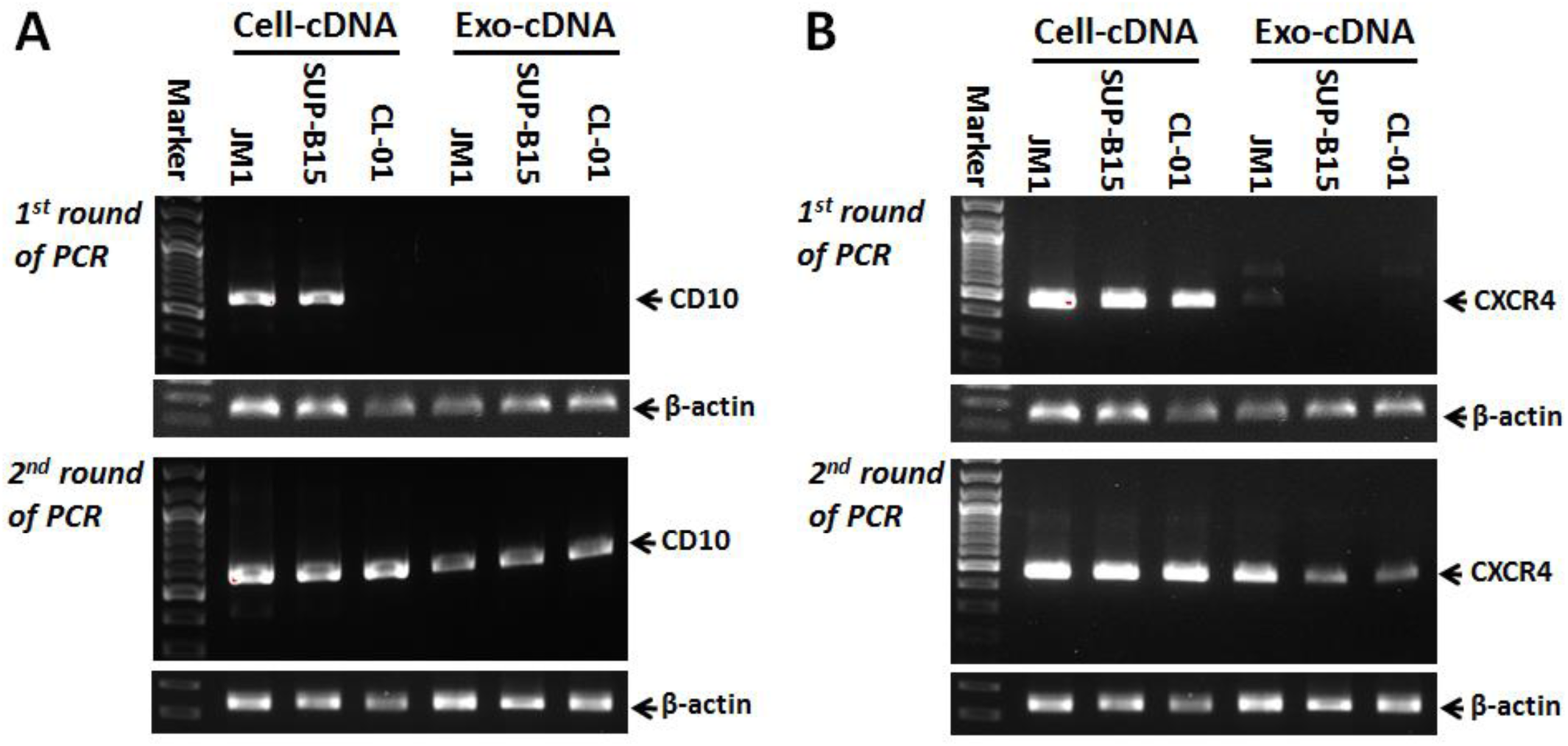
Amplification of cytoplasmic CD10 and CXCR4 from Cell-cDNA and Exo-cDNA. Agarose gel elecprohporesis were carried out after 1st and 2nd round of PCR amplification as shown in upper and lower panel respectively. **A**. CD10 mRNA expression in Cell-cDNA and Exo-cDNA in indicated cell lines (JM1, SUP-B15, and CL-01), upper panel (first round of PCR), lower panel (seocnd round of PCR). **B**. CXCR4 mRNA expression in Cell-cDNA and Exo-cDNA in indicated cell lines (JM1, SUP-B15, and CL-01), upper panel (first round of PCR), lower panel (seocnd round of PCR). β-actin housekeeping gene as quality control.

### 3. 3. Amplication of PCNA and CREB-BP from Cell-cDNA/Exo-cDNA

Both PCNA and CREB-BP are nuclear proteins commonly used as cell markers. PCNA and CREB-BP mRNA were amplified by 1^st^ and 2^nd^ rounds of PCR utilizing Cell-cDNA and Exo-cDNA from each cell lines (JM1, SUP-B15, and CL-01) (**Fig. 3**). In a first round of PCR, PCNA mRNA expression at cellular level (Cell-cDNA) was amplified in all three cell lines but could not amplify at exosomal level (Exo-cDNA) (**Fig. 3 A, upper panel**). Similarly after a second round of PCR, PCNA mRNA expression was detected at cellular level (Cell-cDNA) but exosomal level (Exo-cDNA) did not show PCNA mRNA amplification (**Fig. 3 A, lower panel**). When we tested CREB-BP amplification, a comparable pattern was observed: a first round of PCR showed CREB-BP mRNA amplified product from all three cell lines derived Cell-cDNA, but Exo-cDNA did not produce a detectable level of PCR product (**Fig. 3 B, upper panel**). A second round of PCR demonstrated consistent CREB-BP mRNA amplification of Cell-cDNA, in contrast to Exo-cDNA that again could not amplify a detectable amount of CREB-BP (**Fig. 3 B, lower panel**). Our findings are suggestive of the fact that nuclear mRNA transcripts are not present in the cDNA derived from exosomes. We confirmed this observation by using additional mRNA transcripts that encode other *nuclear* proteins such as PCNA, p53, VBRAF600 and TdT and each time, Exo-cDNA failed to amplify. Endogenous control (β-actin) was used as quality confirmation of Cell-cDNA and Exo-cDNA.

**Figure 3.**
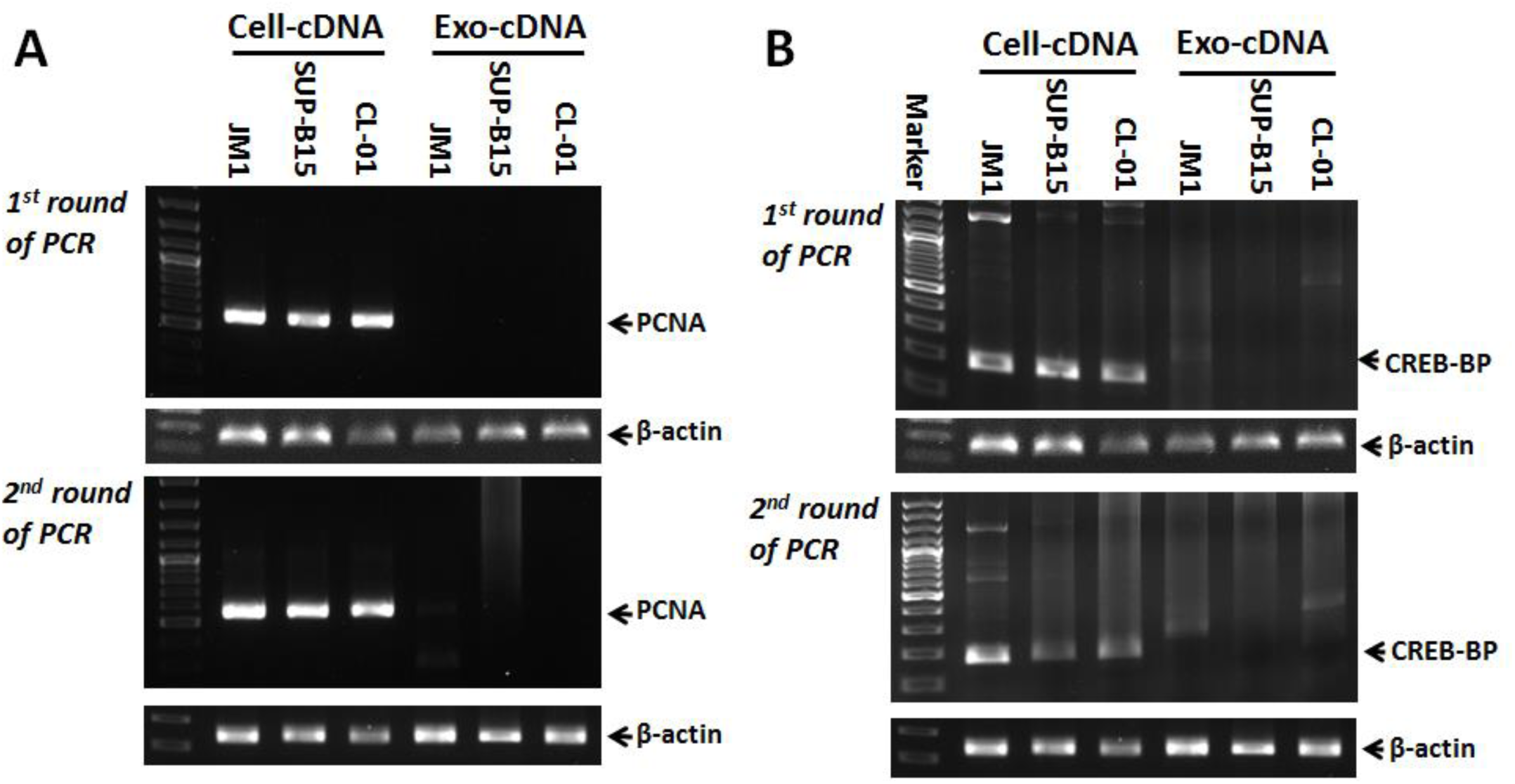
Amplification of nuclear PCNA and CREB-BP from Cell-cDNA and Exo-cDNA. Agarose gel elecprohporesis were carried out after 1st and 2nd round of PCR amplification as shown in upper and lower panel respectively. **A**. PCNA mRNA expression in Cell-cDNA and Exo-cDNA in indicated cell lines (JM1, SUP-B15, and CL-01), upper panel (first round of PCR), lower panel (seocnd round of PCR). **B**. CREB-BP mRNA expression in Cell-cDNA and Exo-cDNA in indicated cell lines (JM1, SUP-B15, and CL-01), upper panel (first round of PCR), lower panel (seocnd round of PCR). β-actin housekeeping gene as quality control.

### 3. 4. Amplication of AID and TdT from Cell-cDNA/Exo-cDNA

We tested two more proteins (AID and TdT) which are enzymatic in nature and expressed predominantly in the nucleus. PCR amplification of AID and TdT mRNA expression was carried out and electrophoresed on an agarose gel (**Fig. 4**). A first round of PCR showed AID mRNA amplification of Cell-cDNA derived from JM1, SUP-B15 and CL-01 cell lines but Exo-cDNA could not produce any AID PCR product and therefore could not be detected on agarose (**Fig. 4 A, upper panel**). The normal B cell line, CL-01, is known to overexpress AID. A second round of PCR demonstrated nuclear protein AID mRNA amplification with Cell-cDNA, but Exo-cDNA could not amplify any detectable amount of AID product (**Fig. 4 A, lower panel**). Similarly, PCR amplification of TdT mRNA encoding this nuclear protein failed to amplify cDNA derived from exosomes (**Fig. 4 B**). CL-01 cells were used as a negative control for TdT expression and housekeeping gene (β-actin) was used as confirmation of Cell-cDNA and Exo-cDNA.

**Figure 4.**
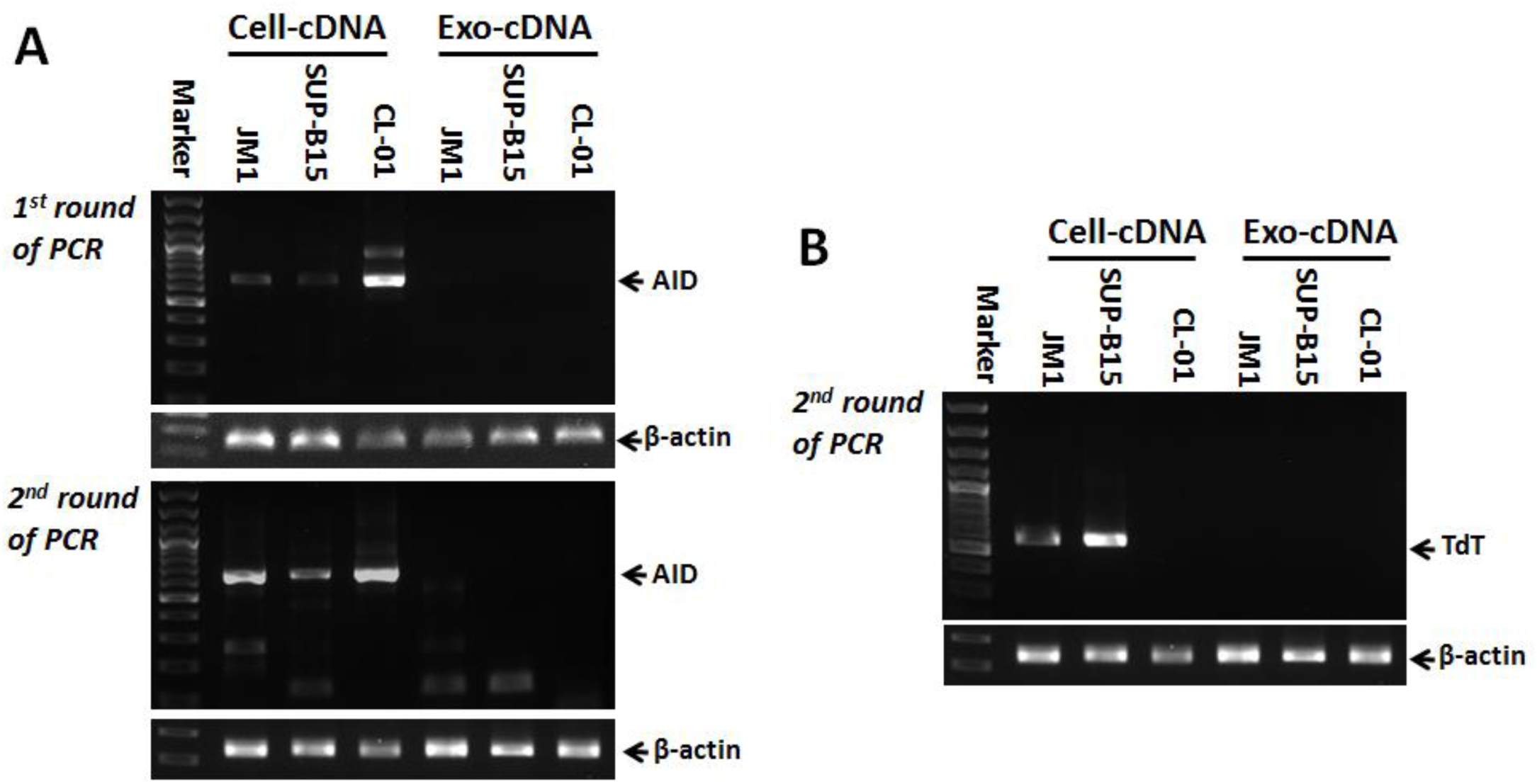
Amplification of nuclear enzymes AID and TdT from Cell-cDNA and Exo-cDNA. Execution of agarose gel elecprohporesis was carried out after 1st and 2nd rounds of PCR amplification as shown in upper and lower panel respectively. **A**. AID mRNA expression in Cell-cDNA and Exo-cDNA in indicated cell lines (JM1, SUP-B15, and CL-01), upper panel (first round of PCR), lower panel (seocnd round of PCR). **B**. TdT mRNA expression in Cell-cDNA and Exo-cDNA in indicated cell lines (JM1, SUP-B15, and CL-01). Represntative data is seocnd round of PCR product. β-actin as housekeeping gene.

Altogether, we have looked at five genes encoding cytoplasmic proteins (CD10, CD34, actin, CXCR4, HLA-DR) and five genes encoding nuclear proteins (p53, PCNA, TdT, CREB-BP, and AID). PCR amplification of all genes are summarized individually as positive and negative in cellular and exosomal cDNA (**Table 2**).

**Table 2:** Total number of cytoplasmic and nuclear coding genes.

| cDNA type | Cell types | Cytoplasmic protein coding genes |  |  |  |  | Nuclear protein coding genes |  |  |  |  |
| --- | --- | --- | --- | --- | --- | --- | --- | --- | --- | --- | --- |
|  |  | CD10 | CD34 | Actin | CXCR4 | HLA-DR | p53 | PCNA | TdT | CREB-BP | AID |
| Cell-cDNA | JM1 | + | + | + | + | + | + | + | + | + | + |
| Cell-cDNA | SUP-B156 | + | + | + | + | + | + | + | + | + | + |
| Cell-cDNA | CL-01 | + | - | + | + | + | + | + | - | + | + |
| Exo-cDNA | JM1 | + | + | + | + | + | - | - | - | - | - |
| Exo-cDNA | SUP-B156 | + | + | + | + | + | - | - | - | - | - |
| Exo-cDNA | CL-01 | + | - | + | + | + | - | - | - | - | - |

## 4. Discussion

Cell biology is defined as the study of the structure and compartments within the cell, while molecular biology can be defined as the study of cellular content within those compartments by analyzing the composition, structure and interaction of cellular molecules such as RNA, protein, DNA etc. The study of biogenesis of exosomes helps us to understand the processing of exosomes by focusing closely on how both fields of biology overlap. This manuscript discusses how the cellular compartmental origin of exosomes delineates and discerns their very specific nature of mRNA transcripts. Our data demonstrate that exosomes encapsulate mRNA transcripts that encode *cytoplasmic* proteins and avoids mRNA transcript encoding *nuclear* proteins.

Exosomes were extracted from the CM of B-cell lines (JM1, SUP-B15, and CL-01) and characterized by NTA system, showing that most of the exosome particle diameter size was 65 nm. Others have reported that a particle size of 30-100 nm in diameter identifies exosomes [14, 15]. We chose to amplify CD10 and CXCR4 mRNA by PCR because both are cytoplasmic proteins. CD10 is expressed by early B, pro-B and pre-B lymphocytes in acute lymphocytic leukemia (ALL) and lymph node germinal centers [16]. CD10 is demonstrated to be cytoplasmic in nature and oriented as a transmembrane region [17]. CD10 is a common ALL antigen and an important cell surface marker in the diagnosis of pediatric ALL. CD10 and CD34 combined expression has prognostic importance and is associated with a better outcome in childhood ALL [18].

CXCR4 is predominantly a cytoplasmic protein but gets shuttle into the nucleus in cancer settings, considered as poor prognostic event [19]. CXCR4 is a well-established biomarker in both solid and liquid tumors. Augmented CXCR4 expression in tumors is associated with low overall survival rate and in general is considered as a poor prognostic feature in cancer [20]. In addition, high expression of CXCR4 as a biomarker in breast cancer, predicts axillary lymph node metastasis [21, 22], and higher risk for bone metastatic disease [23]. CXCR4 along with CXCL12 have been studied as predictive biomarkers of glioma recurrence [24]. Based on studies mentioned, CXCR4 represents a useful prognostic biomarker in cancer [25], CD10 also studied and reported as diagnostic marker in follicular carcinoma (80% patients are positive) and thyroid carcinoma (77%), is not detected in normal thyroid tissue [26]. Based on above reports and evidences we can propose that amplification of exosomal CXCR4 and CD10 has great prospective in early diagnosis or development of diagnostic marker, and prognostic markers in both cancer settings (solid and liquid tumors).

Both acute lymphocytic leukemia cell lines (JM1, and SUP-B15), and the normal B cell line (CL-01) showed amplification of the cytosolic mRNA transcript of CD10 and CXCR4 expression after a second rounds of PCR amplified from both Cell-cDNA and Exo-cDNA. CL-01 has a low expression of CD10, and therefore could not be detected after the first round of PCR.

We chose another set of predominant *nuclear* proteins (PCNA and CREB-BP) for mRNA amplification by two-step PCR. PCNA is a homo-trimer and acts as a scaffold to recruit proteins involved in DNA repair, DNA replication, and chromatin remodeling and epigenetics system [27]. It is a nuclear protein and also an indicator for cell proliferation [28]. CREB-BP is exclusively expressed and localized in the perinuclear and nuclear region [29]. CREB-BP is an ubiquitously expressed gene and mainly participates in the activation of several different transcription factors which helps in embryonic development, growth control, and homeostasis of cells. We found that first and second rounds of PCR showed amplification PCNA and CREB-BP in Cell-cDNA but these two nuclear gene markers could not be detected in Exo-cDNA.

Furthermore, two other nuclear enzymes (AID and TdT) were selected for PCR amplification. Major functions of AID include somatic hypermutation, gene conversion, and class-switch recombination of immunoglobulin genes in B cells of the immune system [30, 31]. AID is highly expressed in germinal center B cells and contributes to the generation of both protective antibodies against microbes and high affinity, class-switched pathogenic auto-antibodies [32]. AID is also a nuclear protein and localized to the sub-nuclear region [33]. There is also evidence that AID shuttles between the nucleus and the cytoplasm like apolipoprotein B [34].

TdT is a unique DNA polymerase enzyme, expressed in a variety of cells such as immature pre-B, pre-T lymphoid cells, ALL and lymphoma cells [16]. TdT catalyzes addition of N-nucleotides to the V, D, and J exons of the TCR and BCR genes during antibody gene recombination which lead to junctional diversity [35]. TdT is a nuclear protein as reported [36]. In our study, we observed that Cell-cDNA of JM1 and SUP-B15 cells amplified AID and TdT in first and second rounds of PCR but again no amplification of AID and TdT was detected when Exo-cDNA was utilized as a template for PCR. Cell-cDNA derived CL-01 cells could not amplify TdT which was expected, since TdT is a marker for malignant B cells and is absent in normal B cells. We did not find mRNA transcripts encoding any of the tested nuclear proteins (TdT, CREB-BP, AID, and PCNA) in the exosomes. These findings intrigued us as to why exosomes are choosing one set of mRNA transcripts and avoiding entrapment of another set of mRNA transcripts during its process of packaging and biogenesis? Structurally, the size of the nucleus is tremendously smaller than the whole cell with the nuclear volume accounting for only 8% of the cellular volume [37]. As a result, cells may have tendencies to conserve mRNA transcripts responsible for encoding nuclear proteins. Moreover, Hocine S and colleagues [10] reported that when mRNA transcripts migrate out of the nucleus after nuclear processing, many proteins which are adhered and tagged on to the mRNA transcripts regulate pathway and fate of the mRNA transcripts. These finding provide a platform to interpret and explain that there may be potentially two separate sets of mRNA transcripts, which are tagged by different sets of proteins directing mRNA transcripts, such that one set of proteins direct mRNA transcripts to enter into the exosomes (mRNA transcripts encoding cytoplasmic proteins) while another set of proteins direct mRNA transcripts to not enter into the exosomes (mRNA transcripts encoding nuclear proteins).

## 5. Conclusion

In conclusion, we show that exosomes carry mRNA transcripts encoding cytoplasmic proteins but not mRNA transcripts encoding nuclear proteins. Since exosomes are widely used as biomarkers for cancer and other diseases, we urge researchers in biomedical medicine to take our observation into account to possibly avoid premature negative conclusions. A selection of exosomal mRNA transcripts encoding *nuclear* proteins for biomarkers studies in exosomes, will not be expressed and therefore might not be the right choice for a biomarker study. In contrast, exosomal RNA transcripts of *cytoplasmic* proteins such as CXCR4/CD10 and others would be our recommended choice as potential exosomal biomarkers in various diseases conditions.

## Acknowledgments

This work was supported by the Pediatric Oncology Fund at Staten Island University Hospital, The DiMartino Family Foundation and Island Auto group. We also thanks Malvern Instruments for the characterization of exosomes by NTA.

## Author’s contribution

S.H. conceived the concept, conducted the experiments, analyzed data and wrote the manuscript. S.V. discussed that data for analysis and reviewed the manuscript.

## Abbreviations

Exo-CM: Exosomes derived from conditioned medium
Exo-cDNA: direct conversion of exosomal RNA into cDNA.

